# Identifiability analysis and noninvasive online estimation of the first-order neural activation dynamics in the brain with closed-loop transcranial magnetic stimulation

**DOI:** 10.1101/2022.07.14.500136

**Authors:** Seyed Mohammad Mahdi Alavi, Adam Mahdi, Fidel Vila-Rodriguez, Stefan M Goetz

## Abstract

**Background:** Neurons demonstrate very distinct nonlinear activation dynamics, influenced by the neuron type, morphology, ion channel expression, and various other factors. The measurement of the activation dynamics can identify the neural target of stimulation and detect deviations, e.g., for diagnosis. This paper describes a tool for closed-loop sequential parameter estimation (SPE) of the activation dynamics through transcranial magnetic stimulation (TMS). The proposed SPE method operates in real time, selects ideal stimulus parameters, detects and processes the response, and concurrently estimates the input–output (IO) curve and the first-order approximation of the activated neural target.

**Objective:** To develop a method for concurrent SPE of the first-order activation dynamics and IO curve with closed-loop TMS.

**Method:** First, identifiability of an integrated model of the first-order neural activation dynamics and IO curve is assessed, demonstrating that at least two IO curves need to be acquired with different pulse widths. Then, a two-stage SPE method is proposed. It estimates the IO curve by using Fisher information matrix (FIM) optimization in the first stage and subsequently estimates the membrane time constant as well as the coupling gain in the second stage. The procedure continues in a sequential manner until a stopping rule is satisfied.

**Results:** The results of 73 simulation cases confirm the satisfactory estimation of the membrane time constant and coupling gain with average absolute relative errors (AREs) of 6.2% and 5.3%, respectively, with an average of 344 pulses (172 pulses for each IO curve or pulse width). The method estimates the IO curves’ lower and upper plateaus, mid-point, and slope with average AREs of 0.2%, 0.7%, 0.9%, and 14.5%, respectively. The conventional time constant estimation method based on the strength-duration (S–D) curve leads to 33.3% ARE, which is 27.0% larger than 6.2% ARE obtained through the proposed real-time FIM-based SPE method in this paper.

**Conclusions:** SPE of the activation dynamics requires acquiring at least two IO curves with different pulse widths, which needs a controllable TMS (cTMS) device with adjustable pulse duration.

**Significance:** The proposed SPE method enhances the cTMS functionality, which can contribute novel insights in research and clinical studies.

## I. Introduction

Transcranial magnetic stimulation (TMS) is a noninvasive method for activating neurons in the brain [1], [2]. It has been approved for diagnosis as well as various therapeutic applications and has become an essential tool in experimental brain research [3]. Most of these applications desire a high stimulation selectivity to activate specific circuits [4]. While the achievable coil focality has physical limits [5], modifications of the pulse shape were found to address differences in the activation dynamics of various neuron populations for stimulation specificity [6]–[13].

Changing the pulse shape does not only allow shifting the activation between different neural elements but also the analysis of the activation dynamics of the stimulated neural target [14]–[18]. Although the neuron activation dynamics is highly nonlinear and therefore complicated to measure, a linear first-order approximation can already disclose differences between targets [10]. As this first-order linearization typically converges to a low-pass filter, the key parameters that can be extracted in such system identification are the time constant and the static gain, also called coupling gain or coupling factor here [17]. The coupling gain depends on the neurostimulation technology including the coil type and geometric conditions, for instance, coil-to-cortex distance, head size, and neural population type as well as orientation relative to the induced electric field in transcranial magnetic stimulation (TMS) [17], [19]. In addition to identifying and differentiating various stimulation targets, the activation dynamics of neurons are furthermore very sensitive to the environment of the neuron, genetic deviations, and diseases. For instance, neurophysiological studies confirm that the time-constant decreases with hyperpolarization and it increases with depolarization [20]. The time constant increases with demyelination and is significantly longer than healthy controls in neurodegenerative disorders such as amyotrophic lateral sclerosis (ALS) and severe spinal muscular atrophy [21]–[23]. After intravenous immunoglobulin (IVIg) therapy, significant reduction of the time constant is seen in chronic inflammatory demyelinating polyneuropathy (CIDP) [24]. Also, sensory fibers represent significantly longer time constant than motor fibers in healthy controls and patients with alcoholic polyneuropathy [25]. Therefore, real-time identification of the neural activation dynamics appears pivotal to the development of the biomarkers for diagnostics and therapeutic applications.

Due to the long pathway between activating primary motor cortex and recording a muscle’s motor evoked potential (MEP), the approximate identification of the activation dynamics through the measurement of the time constant of the activated neural elements is typically performed in targets with readily detectable response, such as the primary motor cortex or rarer the visual cortex [6], [7], [26]–[28]. Significant research is in progress to resolve this limitation by developing new methods which allow electromyographic (EMG) recordings even below the noise floor of background activity, [29]– [32]. However, there approximate model

Conventionally, the time constant is estimated by using the strength–duration (S–D) curve, which is obtained by changing the pulse duration and extracting response-matching stimulation strengths, e.g., the motor threshold, for each of the pulse durations. The time constant is estimated by fitting the Lapicque or Weiss models to the S–D curve [10], [17], [20]. Thus, the S–D-based time constant estimation is off-line, a time consuming process, and records data blindly before they are analyzed.

Real-time estimation of the neural time constant and the coupling factor is a major challenging issue and requires closed-loop stimulation, analysis of the response, and decision about the parameters of the subsequent stimulus [33]. Furthermore, various technologies to change the pulse shape with a sufficiently wide range during operation have been proposed and developed to enable such closed-loop estimation of the time constant [34]–[39].

Closed-loop TMS refers to the automatic and real-time adjustment of TMS parameters based on measurements, e.g., to maximize the desired plastic effects by using the brain/neural data in a feedback system or to speed up the detection of a biomarker [40], [41]. Among the simplest closed-loop TMS procedures are motor threshold estimation methods [42]–[44]. Closed-loop TMS is an area of active research using both EEG-[41], [45]–[48] and EMG-guided TMS [33], [49], [50].

We previously proposed to use real-time sequential parameter estimation (SPE) of the neural membrane time constant and recruitment input–output (IO) curve and developed the necessary analytical relationships for a fixed coupling gain with TMS [33]. In SPE, a number of TMS pulses is administered, the estimation is updated after each stimulus, and the estimation continues until satisfying a stopping rule based on the convergence of the model parameters [49].

As the key objective, this paper presents a tool for concurrent SPE of the membrane time constant and IO curve parameters including the coupling gain with closed-loop EMG-guided TMS. To achieve this goal, an integrated model of the full first-order neural membrane dynamics and IO curve is firstly developed. The identifiability analysis demonstrates that at least two IO curves with different pulse widths are required for concurrent sequential parameter estimation of the membrane time constant and coupling gain. This emphasizes the need for controllable TMS (cTMS) devices with adjustable pulse width. Then, a two-stage method is proposed for SPE of the IO curve in the first stage and the membrane time constant as well as the coupling gain in the second stage. The IO curve is estimated by optimizing the Fisher information matrix (FIM) [49]. The results are validated through extensive simulations and compared with the conventional time-constant estimation using the S–D curve. The proposed method enhances the methodology and functionality of TMS and promises novel insights in the physiology and clinical applications.

### A. The contributions of this paper

The contributions of this paper are highlighted as follows:

– Formal identifiability analysis of the integrated model of the full first-order neural membrane dynamics and IO curve, which underlines that at least two IO curves at different pulse widths need to be acquired for concurrent sequential parameter estimation of the membrane time constant and coupling gain.
– Concurrent sequential parameter estimation of the full first-order dynamical model of the neural membrane (including the time constant and coupling gain) and pulse-width-dependent IO curve with closed-loop EMG-guided cTMS.
– Comparison between the proposed SPE and S–D curve based time constant estimation methods.

### B. The structure of the paper

This paper is organized as follows: Section II explains an integrated model of the neural activation dynamics and IO curve. Section III states the problem from the mathematical perspective. Section IV discusses the identifiability of the model parameters. The proposed sequential parameter estimation method is described in Section V, and finally, Section VI discusses the simulation method, results, and a comparative study.

## II. Neural system model

Fig. 1 shows the overall structure of the neural system model, from the TMS pulse *w* to the MEP size *y*. The MEP size can be represented by various metrics, such as peak-to-peak voltage, area, or similar features [31]. *h*(*t*) represents the dynamics of the neural membrane. This paper focuses on the first-order linear approximation model with the impulse-response function

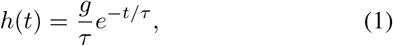

where *τ* is the membrane time constant and *g* is the coupling gain between the TMS coil and the directly stimulated neurons, which can depend on factors such as the coil including shape and number of turns, coil-to-cortex distance and specific coil position, anatomy, properties of the neurons in the focus, including morphology and ion-channel expression, and orientation relative to the induced electric field [17], [18].

**Fig. 1.**
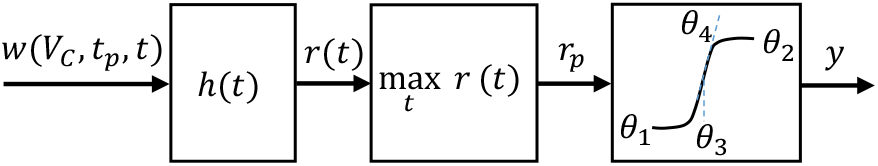
The overall structure of the neural system model, from the TMS pulse *w* to the MEP size *y*.

*r*(*t*) denotes the membrane response to the stimulus *w*. Any TMS technology that can vary the pulse duration sufficiently can serve for this time-constant determination. Without loss of generality, this paper uses the original cTMS design [35] as an example with

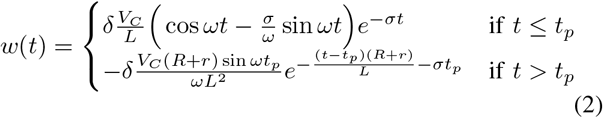

where *V*_*C*_ is the normalized pulse voltage amplitude, adjustable between *V*_*C*_(min) = 0 and *V*_*C*_(max) = 1. *t*_*p*_ is the pulse width, which is assumed to be adjustable between *t*_*p*_(min) = 10 *µ*s and *t*_*p*_(max) = 200 *µ*s. Other parameters include the stimulating coil *L*, the energy dissipation resistor *R* of the free-wheeling path that generates the decay of the second pulse phase, the resistor *r* which represents the combined resistance of the capacitor, semiconductor switch, cables, and stimulation coil. In this paper, the following parameter values are used based on the original cTMS design [35]: *L* = 16 *µ*H, *C* = 716 *µ*F, *R* = 0.1 Ω, *r* = 20 mΩ, *δ* = 3.2 × 10^−6^ (V/m)(A/s). The parameters *ω* and *σ* are defined as

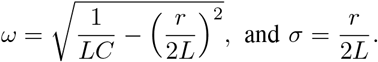

The parameter *δ* is a proportionality coefficient.

The membrane response *r*(*t*) is obtained by the convolution of *w* and *h* as follows [33]

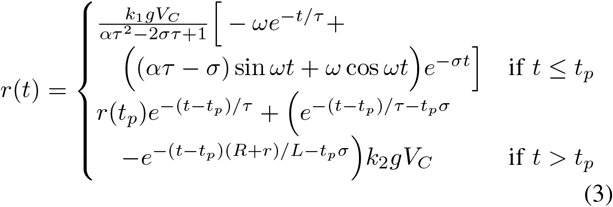

where *r*(*t*_*p*_) is the neural membrane response at time *t* = *t*_*p*_, and

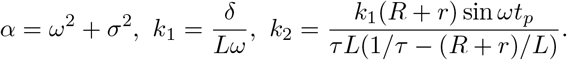

In Fig. 1, *r*_*p*_ is called the depolarization factor, which points out the peak of *r* as follows

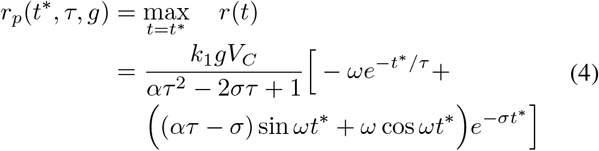

where *t*\* is the peak time.

An MEP is generated when the depolarization factor *r*_*p*_ reaches a certain threshold. Thus, the relationship between the depolarization factor and the MEP is modeled as a sigmoid function

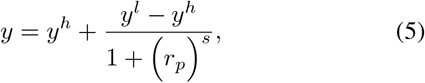

where *y* is the MEP size and *y*^*l*^ as well as *y*^*h*^ represent the lower and upper plateaus of the IO curve.

By defining the parameter vector

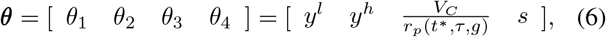

and considering the neurophysiological and technical uncertainties and variabilities [51], [52], an integrated model of the TMS neural system is obtained as

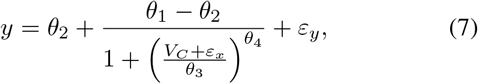

where *ε*_*x*_ and *ε*_*y*_ represent the neurophysiological and technical uncertainties and variabilities such as trial-to-trial variabilities, excitability fluctuations, variability in the neural and muscular pathways, physiological and measurement noise, and parameter uncertainties.

The mid point and slope of the IO curve are given by *θ*_3_ and *θ*_4_, respectively.

Fig. 2 presents a sample cTMS pulse *w*(*t*), the first-order linear neural response *r*(*t*), and the depolarization factor *r*_*p*_.

**Fig. 2.**
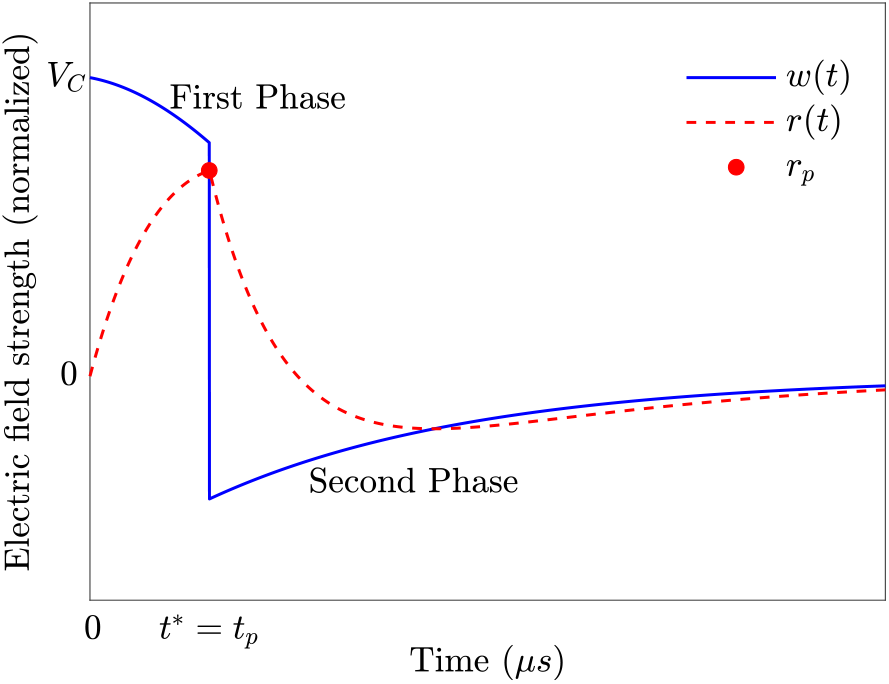
Sample cTMS pulse *w*(*t*), first-order linear neural response *r*(*t*), and depolarization factor *r*_*p*_.

## III. Problem Statement

The problem is to estimate the integrated model parameters, i.e., neural membrane time constant *τ*, gain *g*, and the IO curve parameters *θ*_*i*_, *i* = 1, …, 4 with closed-loop cTMS. Most importantly, all those parameters need to be estimated concurrently. The adjustable parameters are the pulse amplitude *V*_*C*_ and the pulse width *t*_*p*_. This paper will discuss the identifiability conditions for such a concurrent estimation to subsequently provide a method.

## IV. Identifiability analysis

The proposed method is based on a two-stage SPE technique, where the estimation of the parameter vector ***θ*** is firstly updated after each stimulus. Then, the membrane time constant and coupling gain estimations are updated by using the relationship between the IO curve’s mid-point *θ*_3_ and these parameters as follows

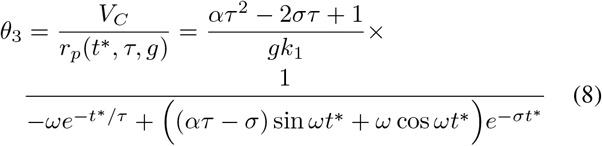

where, *t*\* is the peak time. For the given TMS pulse (2), if the pulse width is shorter than a critical pulse width, 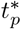, the peak time will occur at the end of the pulse. Otherwise, *t*\* occurs before that, as discussed in [33], [53]. The relationship between the time constant and pulse width is given by

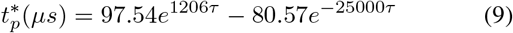

For identifiability, it is then required to assume

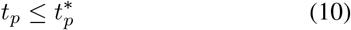

which results in

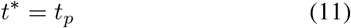

There are two unknown parameters *g* and *τ* on the right side of (8). Thus, it is required to acquire at least two IO curves at different pulse widths to estimate *τ* and *g*. Since stimulation at two pulse widths is in principle sufficient for the estimation of these parameters, this paper focuses on two IO curves. In this case, Eq. (8) is re-written as

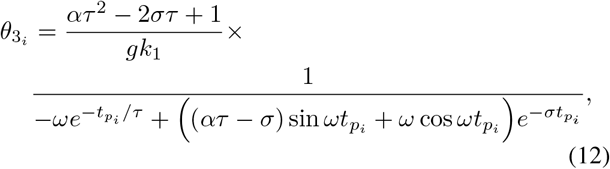

where 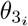 and 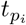 are the mid point and pulse width of the *i*−th IO curve with *i* = 1, 2.

The membrane time constant *τ* is obtained by solving 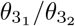 as follows:

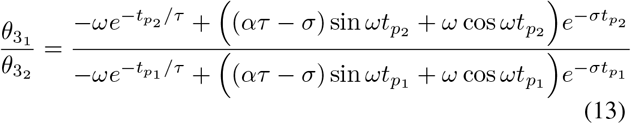

The gain *g* is computed by solving (12) for *i* = 1 or 2.

## V. Sequential parameter estimation

As discussed earlier, at least two IO curves with different pulse widths 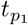 and 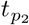 are required. Based on (10), 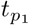 and 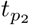 are chosen arbitrarily between *t*_*p*_(min) and 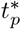. Denote the parameter vectors of the first and second IO curves separately as

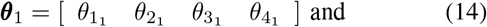

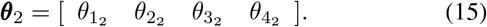

The SPE method starts by taking samples from the baseline.

Subsequently, the pulse width of the cTMS device is set to 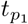, three initial pulse amplitudes are chosen randomly between *V*_*C*_(min) and *V*_*C*_(max), and the corresponding MEPs are measured. Then, the pulse width of the cTMS device is set to 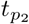, three initial pulse amplitudes are chosen randomly between *V*_*C*_(min) and *V*_*C*_(max), and the corresponding MEPs are measured. The initial stimuli could be performed at the same pulse amplitudes for both pulse widths.

In the next step, initial estimations of ***θ***_1_ and ***θ***_2_ are obtained by fitting the sigmoid model (7) to the baseline and initial stimuli-response data. The membrane time constant and gain *g* are then computed by using the estimated 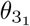 and 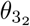 as described in the previous section.

The next pulse amplitudes, i.e., 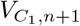 for the IO curve estimation with the pulse width 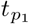 and 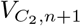 for the IO curve estimation with the pulse width 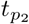, are chosen and administered by using the prior data 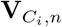 and 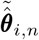, and optimizing the Fisher information matrix (FIM) in a closed-loop feedback system as follows

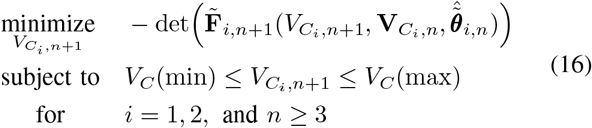

with

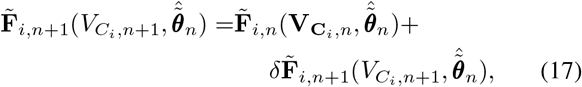

where 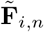 is FIM at *n*−th stimulus, 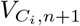 is the next pulse amplitude, 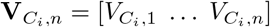 is the pulse amplitude’s prior data, and 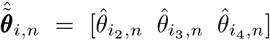 is the parameter vector’s prior data for the *i*−th IO curve.

Although the FIM optimization (16) assumes normal distributions for the variabilities, simulation studies illustrate its robustness to variabilities with asymmetric distributions. Further details have been discussed previously [49].

The MEP characteristics are measured, and the data sets are updated for both IO curves. The estimations of ***θ***_1_ and ***θ***_2_ are updated by fitting the sigmoid model (7) to the baseline and updated stimuli-response data. The estimation of *τ* and *g* are then updated by using the most recent estimates of 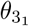 and 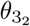 as described in the previous section. This process is continued until the convergence criterion

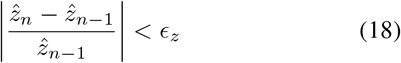

is satisfied for *T* consecutive times, *T* ≥ 1, for all parameters 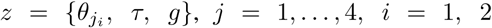. 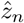 denotes the estimate of *z* after the *n*–th stimulus. *ϵ*_*z*_ denotes the convergence tolerance. The estimation accuracy is adjustable by *ϵ*_*z*_ and *T* values. The larger the *T* value and the smaller the *ϵ*_*z*_ values, the more the accurate estimation is obtained at the cost of more pulses, [49].

## VI. Simulation Results

The effectiveness of the proposed sequential parameter estimation method is evaluated through 73 simulation runs in Matlab R2021a (The MathWorks, Inc.).

In each run, a true membrane dynamics *h*(*t*) is generated with the time constant randomly chosen between 90 and 220 *µ*s. Two pulse widths 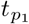 and 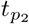 are randomly selected between 10 *µ*s and 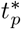, which is computed through (9). In practice, because there is no prior approximate information about the neural membrane time constant, it is suggested to choose short pulse widths.

By randomly choosing the coupling gain *g* between 30 and 50, the true values of the IO curve’s mid points 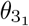 and 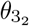 are computed, which are real scalars between 0 and 1. Without loss of generality, the lower and upper plateaus are assumed to be the same for both IO curves, i.e., 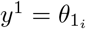, and 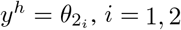. The true value of *y*^*l*^ is randomly chosen between –6.5 and –5.5 (corresponding to 0.32 and 3.2 *µ*V), and the true value of *y*^*h*^ is randomly selected between –3 and –2 (corresponding to 1 and 10 mV). True values of the slopes, 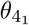 and 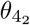, are randomly chosen between 1 and 100. The reference IO curves are generated by using these random data for all 73 runs. The stimulus–response pairs are generated by applying Gaussian noise with standard deviations of 0.05 and 0.1 to the x and y axes, respectively [51], [54].

The problem in this section is to concurrently estimate the true values of the membrane time constant, coupling gain, and IO curves’ parameters by using the proposed two-stage FIM-based SPE method.

Fifty baseline samples are arbitrarily taken for all IO curves in all runs.

Every time the estimation of the IO curves and parameter vectors is updated, the trust-region curve fitting algorithm is run with lower and upper limits on the parameter vector as follows:

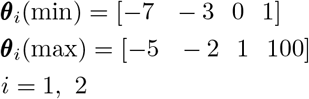

The curve fitting is performed on the logarithmic scale to mitigate the highly skewed variabilities’ effects [10], [17]. A bad-fit detection and removal technique is used to avoid sudden estimation jumps.

The estimation of the time constant is updated by solving (13) using fmincon and the global search interior-point algorithm with a random initial guess, and lower and upper limits of 90 *µ*s and 220 *µ*s, respectively.

The optimization problem (16) is solved by using fmincon and the global search interior-point algorithm with a random initial guess, and lower and upper limits of 0.01 and 1, respectively.

For each IO curve, the maximum number of pulses is arbitrarily set to *n*_max_ = 500, which means that 1000 stimuli can be administered in total.

The stopping rule is based on *T* = 5 successive satisfaction of the the convergence criterion (18) with the tolerance *ϵ*_*z*_ set to 0.01 for all parameters.

### A. The results of a representative run

Fig. 3 shows the stimulus–response pairs (‘×’ signs) and reference IO curves (solid lines) for a representative run with the following arbitrarily chosen true values:

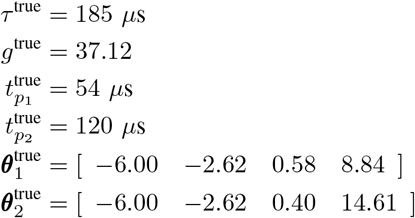

**Fig. 3.**
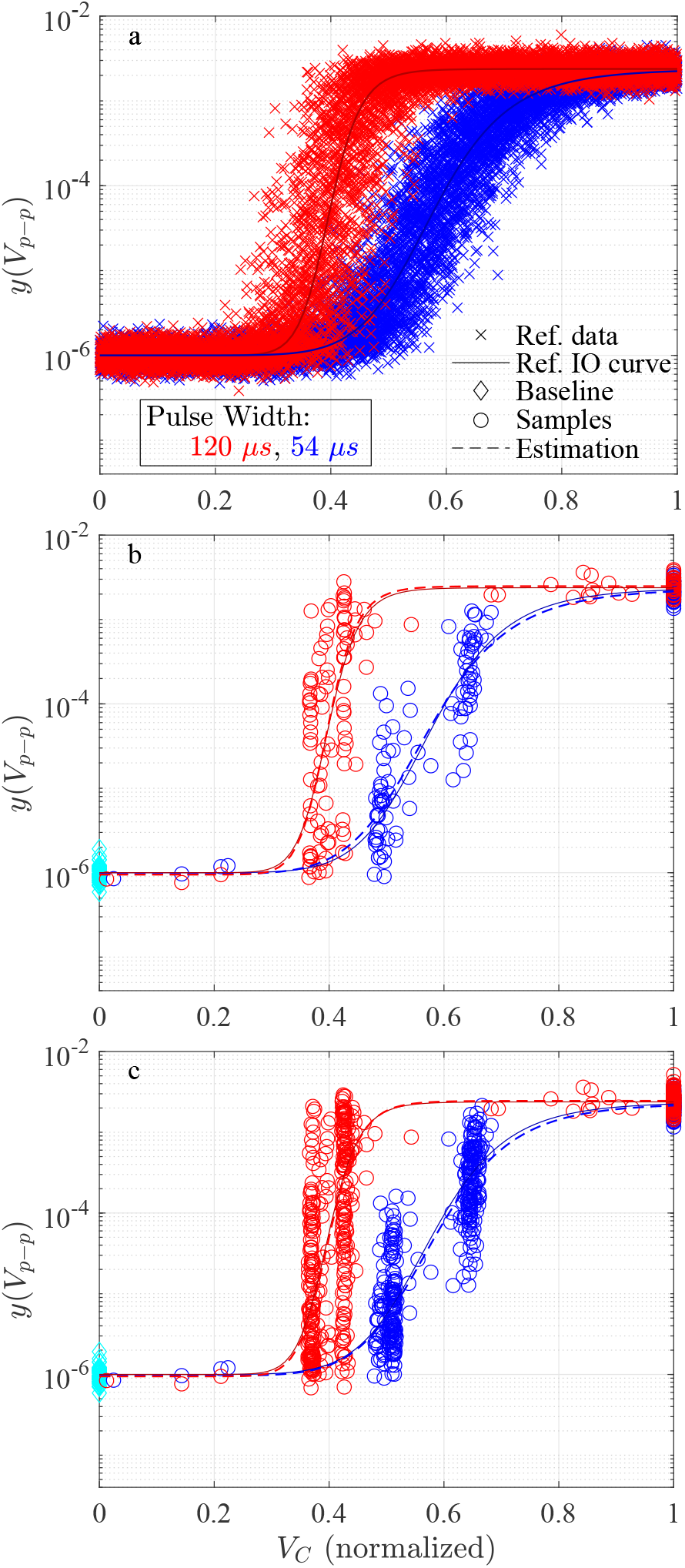
Sample simulation run: (a) reference data and IO curves at 54 and 120 *µs*. Estimation of the IO curves by using the proposed method at (b) *n* = *n*_*f*_ = 149, when the stopping rule is satisfied, and (c) *n* = *n*_max_ = 500, when the maximum number of pulses is administered. It is seen that the FIM-based IO-curve estimation dominantly administers pulses from the points of maximum information, which are at *V*_*C*_ (max) and in two areas on the slope.

The critical pulse width 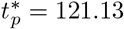 *µ*s is obtained by solving (9) for *τ*^true^ = 185 *µ*s. As discussed in the identifiability section IV, the pulse widths should be shorter than 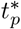.The sequential parameter estimation method satisfies the stopping rule at *n* = *n*_*f*_ = 149 for each IO curve, for this representative run. This means that the parameters have been estimated after administering 2 × 149 = 298 stimuli.

Fig. 3-b and Fig. 3-c summarize the stimulus–response pairs as well as the estimated IO curves at *n*_*f*_ = 149 and *n*_max_ for this case study. As discussed in [49], it is seen that the FIM optimization administers stimuli, mainly from sectors which contain the maximum information for curve fitting. Fig. 4 shows the estimations of the IO curves’ parameters versus the administered pulses. Fig. 5 and Fig. 6 present the estimated time constant and coupling gain, respectively. The results confirm satisfactory estimation of the membrane time constant. coupling gain, IO curves and their parameters.

**Fig. 4.**
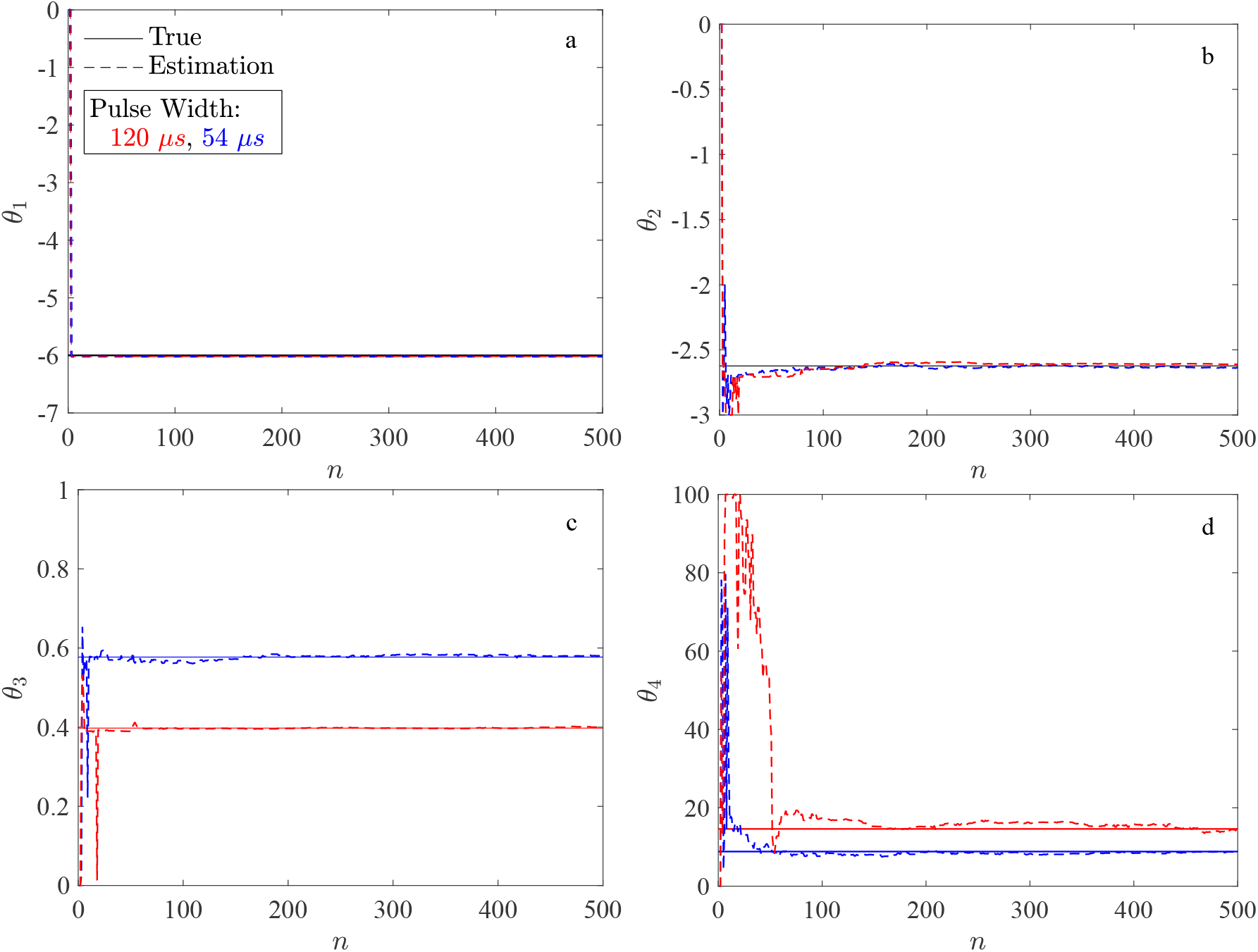
Estimation of the IO curves parameters versus TMS pulses, and comparison with the true values. a) The lower plateau *θ*_1_. b) The upper plateau *θ*_2_. c) The mid-point *θ*_3_. d) The slope *θ*_4_.

**Fig. 5.**
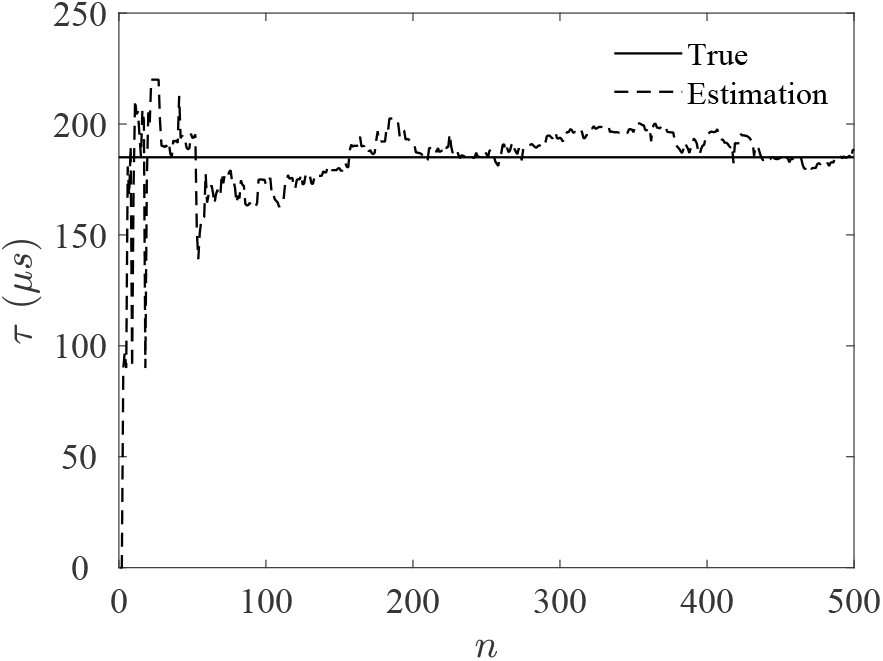
Estimation of the membrane time constant *τ* versus TMS pulses, and comparison with the true value.

**Fig. 6.**
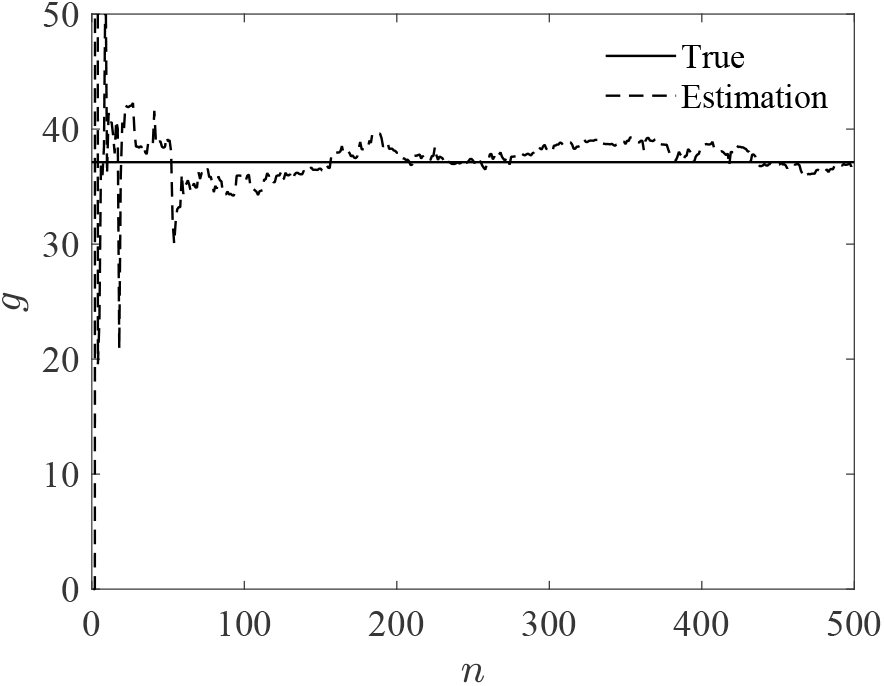
Estimation of the coupling gain *g* versus TMS pulses, and comparison with the true value.

### B. The results of 73 runs

All 73 runs satisfied the stopping rule with an average of *n*_*f*_ = 172 stimuli for each IO curve, which means that 2 × 172 = 344 stimuli were administered in average for the estimation of all parameters and the IO curves.

For all parameters, the absolute relative estimation errors (AREs) after the stimulation of the *n*-th stimulus for each IO curve are computed per

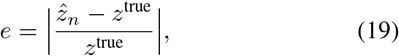

where 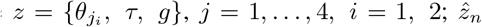 denotes the estimation of *z* after the *n*-th stimulus.

When the stopping rule is satisfied, the IO curves’ parameters *θ*_1_, *θ*_2_, *θ*_3_, and *θ*_4_ are estimated with the average AREs of 0.2%, 0.7%, 0.9%, and 14.5%, respectively. The membrane time constant *τ* and gain *g* are estimated with the average AREs of 6.2% and 5.3%, respectively.

At *n* = *n*_max_ = 500, the IO curves’ parameters *θ*_1_, *θ*_2_, *θ*_3_, and *θ*_4_ are estimated with the average AREs of 0.2%, 0.4%, 0.6%, and 8.7%, respectively. The membrane time constant *τ* and gain *g* are estimated with the average AREs of 5.2% and 4.3%, respectively, at *n* = *n*_max_.

It is noted that the estimation performance at *n* = *n*_*f*_ can be controlled through the convergence tolerance *ϵ*_*z*_ and the number of successive times the convergence criteria must be satisfied; for example, it could be improved by reducing the former or increasing the latter at the cost of a longer procedure needing more TMS pulses.

### C. Comparison with SD-curve-based time constant estimation

Conventionally, the neural time constant *τ* is obtained by fitting the Weiss model [55], [56]

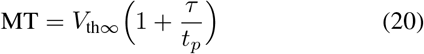

off-line, i.e., in the aftermath, to strength–duration (S–D) data, e.g., motor threshold MT levels at various different pulse widths *t*_*p*_. The parameter *V*_th∞_ is the rheobase amplitude, which implies the smallest pulse amplitude required for threshold stimulation for an infinitely long stimulus pulse.

In order to describe the comparison method, consider the representative run discussed above. Two IO curves are identified at two different pulse widths. The MT associated with the *i*-th IO curve, MT_*i*_, is obtained by solving

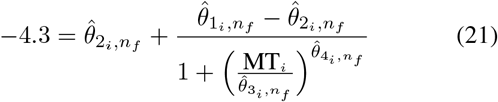

where, 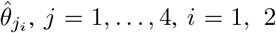, denotes the estimation of 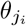 when sequential parameter estimation method successfully stops at *n* = *n*_*f*_. Due to the fitting in the logarithmic scale, log_10_(50 × 10^−6^) = −4.3 is used. The S–D curve time constant is obtained by fitting the Weiss model (20) to the data set 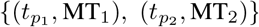. For the representative run, MT_1_ = 57.44% and MT_2_ = 39.71%. By using the same curve fitting method discussed above, the time constant is estimated as 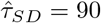 *µs* with the relative error of 51.35%.

The same comparison method over all 73 runs leads to 33.29% ARE at *n* = *n*_*f*_ for the SD-curve-based time constant estimation, which is 27.01% larger than 6.2% ARE obtained through the proposed real-time FIM-based SPE method in this paper.

The result of the SD-curve-based time constant estimation could be improved by capturing more MTs at more pulse width. For instance, by capturing two additional MTs, 93.07% and 48.53%, associated with the pulse widths 30 *µs* and 70 *µs*, the time constant is estimated as 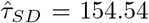 *µs* with the ARE of 16.46%. Fig. 7 illustrates the corresponding estimated SD curves with two and four pulse widths.

**Fig. 7.**
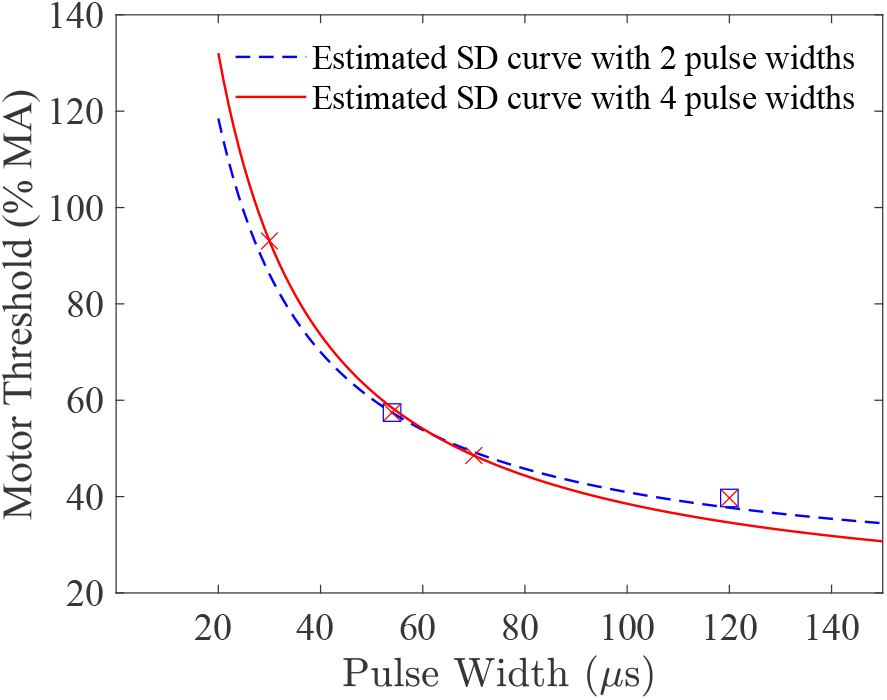
Estimation of strength-duration (S–D) curves with two and four pulse widths. The MT values are obtained after estimation of the IO curve parameters, by solving Eq. (21).

Thus, compared to the SD-based estimation, the proposed method in this paper estimates the time constant with far better accuracy and fewer pulses.

## VII. Conclusions and future works

TMS devices with adjustable pulse duration, such as cTMS, allow closed-loop sequential parameter estimation (SPE) of the linearized neural membrane time constant, the input–output (IO) curve, as well as the coupling gain. It was shown that stimulation at two pulse widths is in principle sufficient for the estimation of these parameters with electromyography-guided (EMG-guided) closed-loop TMS. The proposed SPE method is a two-stage method, estimating the IO curve parameters by using the Fisher information (FIM) matrix optimization in the first stage and estimating the neural time constant and coupling gain in the second stage. The validation with 73 realistic simulation case studies demonstrated satisfactory average absolute relative estimation errors (AREs), specifically 6.2% and 5.3% for the time constant and coupling gain, and 0.2%, 0.7%, 0.9%, and 14.5% for the IO curve parameters, with on average 344 pulses. The ARE of time constant is reduced by 27.09% compared to the conventional off-line method using the strength–duration (S–D) curve. Robustness of the proposed method, increasing the pulse width over the critical pulse width, and a practical easily usable implementation of the proposed method remain for future work. Furthermore, the identification of higher order linear as well as nonlinear components of the activation dynamics may allow further insights into the neurophysiology for instance for the selective stimulation, i.e., the separation of different activated neuron populations.

## References

[1] Polson MJ, Barker AT, Freeston IL. Stimulation of nerve trunks with time-varying magnetic fields. Medical & biological engineering & computing 1982; 20(2):243–244.

[2] Barker AT, Jalinous R, Freeston IL. Non-invasive magnetic stimulation of human motor cortex. The Lancet 1985; 325(8437):1106–1107.

[3] Ziemann U. Thirty years of transcranial magnetic stimulation: where do we stand? Experimental Brain Research 2017; 235:973-–984.

[4] Valero-Cabré A, Amengual JL, Stengel C, Pascual-Leone A, and Coubard OA, Transcranial magnetic stimulation in basic and clinical neuroscience: A comprehensive review of fundamental principles and novel insights. Neuroscience & Biobehavioral Reviews 2017; 83:381–404.

[5] Gomez LJ, Goetz SM, Peterchev AV. Design of transcranial magnetic stimulation coils with optimal trade-off between depth, focality, and energy. Journal of Neural Engineering 2018; 15:046033.

[6] Kammer T, Beck S, Erb M, Grodd W. The influence of current direction on phosphene thresholds evoked by transcranial magnetic stimulation. Clinical Neurophysiology. 2001; 112(11):2015–2021.

[7] Kammer T, Beck S, Thielscher A, Laubis-Herrmann U, Topka H. Motor thresholds in humans: a transcranial magnetic stimulation study comparing different pulse waveforms, current directions and stimulator types. Clinical Neurophysiology. 2001; 112(2):250–258.

[8] Sommer M, Alfaro A, Rummel M, Speck S, Lang N, Tings T, Paulus W. Half sine, monophasic and biphasic transcranial magnetic stimulation of the human motor cortex. Clinical Neurophysiology. 2006; 117(4):838–844.

[9] Goetz SM, Luber B, Lisanby SH, Murphy DLK, Kozyrkov IC, Grill WM, Peterchev AV. Enhancement of neuromodulation with novel pulse shapes generated by controllable pulse parameter transcranial magnetic stimulation. Brain Stimulation. 2016; 9(1):39–47.

[10] D’Ostilio K, Goetz SM, Hannah R, Ciocca M, Chieffo R, Chen J-C A, Peterchev AV, Rothwell JC. Effect of coil orientation on strength–duration time constant and I-wave activation with controllable pulse parameter transcranial magnetic stimulation. Clinical Neurophysiology 2016; 127:675–683.

[11] Sommer M, Ciocca M, Chieffo R, Hammond P, Neef A, Paulus W, Rothwell JC, Hannah R. TMS of primary motor cortex with a biphasic pulse activates two independent sets of excitable neurones. Brain Stimulation. 2018; 11(3):558–565.

[12] Halawa I, Shirota Y, Neef A, Sommer M, Paulus W. Neuronal tuning: selective targeting of neuronal populations via manipulation of pulse width and directionality. Brain Stimulation. 2019; 12(5):1244–1252.

[13] Hannah R, Rothwell JC. Pulse Duration as Well as Current Direction Determines the Specificity of Transcranial Magnetic Stimulation of Motor Cortex during Contraction. Brain Stimulation 2017; 10:106—115.

[14] Barker AT, Garnham CW, Freeston IL. Magnetic nerve stimulation: the effect of waveform on efficiency, determination of neural membrane time constants and the measurement of stimulator output. Electroencephalogr Clin Neurophysiol Suppl 1991; 43:227—37.

[15] Corthout E, Barker AT, Cowey A. Transcranial magnetic stimulation. Which part of the current waveform causes the stimulation? Exp Brain Res 2001; 141:128-–32.

[16] Amassian VE, Maccabee PJ. Transcranial Magnetic Stimulation. International Conference of the IEEE Engineering in Medicine and Biology Society (EMBS) 2006; 22:1620–1623. doi: 10.1109/IEMBS.2006.259398.

[17] Peterchev AV, Goetz SM, Westin GG, Luber B, Lisanby SH. Pulse width dependence of motor threshold and input–output curve characterized with controllable pulse parameter transcranial magnetic stimulation. Clinical Neurophysiology 2013; 124:1364–1372.

[18] Aberra AS, Peterchev AV, Grill WM. Biophysically realistic neuron models for simulation of cortical stimulation. Journal of neural engineering. 2018; 15(6):066023.

[19] Goetz SM, Weyh T, Afinowi IAA, Herzog HG. Coil Design for Neuromuscular Magnetic Stimulation Based on a Detailed 3-D Thigh Model. IEEE Transactions on Magnetics 2014; 50(6):1–10.

[20] Mogyoros I, Kiernan MC, Burke D. Strength-duration properties of human peripheral nerve. Brain 1996; 119 (Pt 2):439–47.

[21] Farrar MA, Vucic S, Lin CS, Park SB, Johnston HM, du Sart D, et al. Dysfunction of axonal membrane conductances in adolescents and young adults with spinal muscular atrophy. Brain. 2011; 134:3185–3197.

[22] Moldovan M, Alvarez S, Pinchenko V, Marklund S, Graffmo KS, Krarup C. Nerve excitability changes related to axonal degeneration in amyotrophic lateral sclerosis: Insights from the transgenic SOD1(G127X) mouse model. Exp Neurol. 2012; 233:408–420.

[23] Lugg A, Schindle M, Sivak A, Tankisi H, Jones KE. Nerve excitability as a biomarker for amyotrophic lateral sclerosis: a systematic review and meta-analysis. medRxiv 2022; doi: https://doi.org/10.1101/2022.02.11.22270866

[24] Lin CS, Krishnan AV, Park SB, Kiernan MC. Modulatory effects on axonal function after intravenous immunoglobulin therapy in chronic inflammatory demyelinating polyneuropathy. Arch Neurol. 2011; 68:862–869.

[25] Yerdelen D, Koc F, Uysal H. Strength–duration properties of sensory and motor axons in alcoholic polyneuropathy. Neurological Research 2008; 30(7): 746–750.

[26] Rossini PM. Methodological and physiological aspects of motor evoked potentials. Electroencephalography and Clinical Neurophysiology Suppl. 1990; 41:124–133.

[27] Ghaly RF, Stone JL, Aldrete JA, and Kartha RK. Transcranial magnetic induced motor evoked potentials in primates: the technique and anesthetic effects. Images of the Twenty-First Century. IEEE Proceedings of the Annual International Engineering in Medicine and Biology Society, 1989; 1573–1574.

[28] Siebner HR, Funke K, Aberra AS, Antal A, Bestmann S, Chen R, Classen J, Davare M, Di Lazzaro V, Fox PT, Hallett M, Karabanov AN, Kesselheim J, Beck MM, Koch G, Liebetanz D, Meunier S, Miniussi C, Paulus W, Peterchev AV, Popa T, Ridding MC, Thielscher A, Ziemann U, Rothwell JC, Ugawa Y. Transcranial magnetic stimulation of the brain: What is stimulated? - A consensus and critical position paper. Clin Neurophysiol. 2022 Aug;140:59–97.

[29] Klomjai W, Katz R Lackmy-Vallée. Basic principles of transcranial magnetic stimulation (TMS) and repetitive TMS (rTMS). Annals of Physical and Rehabilitation Medicine 2015; 58(4):208–213.

[30] Westin GG, Bassi BD, Lisanby SH, Luber B. Determination of motor threshold using visual observation overestimates transcranial magnetic stimulation dosage: safety implications. Clinical Neurophysiology 2014; 125(1):142–147.

[31] Li Z, Peterchev AV, Rothwell JC, Goetz SM. Detection of Motor-Evoked Potentials Below the Noise Floor: Rethinking the Motor Threshold. bioRxiv 2021.12.10.472126. doi: 10.1101/2021.12.10.472126

[32] Goetz SM, Li Z, Peterchev AV. Noninvasive detection of motor-evoked potentials in response to brain stimulation below the noise floor—how weak can a stimulus be and still stimulate. Proc IEEE Eng Med Biol Soc (EMBC) 2018; 40:2687–2690. doi: 10.1109/EMBC.2018.8512765

[33] Alavi SMM, Vila-Rodriguez F, Mahdi A, Goetz SM. A formalism for sequential estimation of neural membrane time constant and input–output curve towards selective and closed-loop transcranial magnetic stimulation. Journal of Neural Engineering 2022; 19(5):056017.

[34] Goetz SM and Deng ZD. The development and modelling of devices and paradigms for transcranial magnetic stimulation. International Review of Psychiatry 2017; 29(2):115–145.

[35] Peterchev AV, Jalinous R, Lisanby SH. A Transcranial Magnetic Stimulator Inducing Near-Rectangular Pulses With Controllable Pulse Width (cTMS). IEEE Trans Biomed Eng 2008;55(1):257–66.

[36] Goetz SM, Pfaeffl M, Huber J, Singer M, Marquardt R, Weyh T. Circuit topology and control principle for a first magnetic stimulator with fully controllable waveform. Proc IEEE Eng Med Biol Soc (EMBC) 2012; 34:4700–4703. doi: 10.1109/EMBC.2012.6347016

[37] Li Z, Zhang J, Peterchev AV, Goetz SM. Modular Pulse Synthesizer for Transcranial Magnetic Stimulation with Flexible User-Defined Pulse Shaping and Rapidly Changing Pulses in Sequences. 2022; 2202.06530. doi: 10.48550/ARXIV.2202.06530

[38] Zeng Z, Koponen LM, Hamdan R, Li Z, Goetz SM, Peterchev AV. Modular multilevel TMS device with wide output range and ultrabrief pulse capability for sound reduction. Journal of Neural Engineering. 2022; 19(2):026008.

[39] Polson MJR. Magnetic stimulator for neuro-muscular tissue. US Patent and Trademark Office 1998; US 5, 766,124.

[40] Karabanov A, Thielscher A, Siebner HR. Transcranial brain stimulation: Closing the loop between brain and stimulation. Curr Opin Neurol 2016;29(4):397-–404.

[41] Tervo AE, Nieminen JO, Lioumis P, Metsomaa J, Souza VH, Sinisalo H, Stenroos M, Sarvas J, Ilmoniemia RJ. Closed-loop optimization of transcranial magnetic stimulation with electroencephalography feedback, Brain Stimulation 2021; 14:P1674.

[42] Wang B, Peterchev AV, and Goetz SM. Analysis and Comparison of Methods for Determining Transcranial Magnetic Stimulation Thresholds. bioRxiv 2022; 495134v1. doi: 10.1101/2022.06.26.495134

[43] Treutwein B and Strasburger H. Fitting the psychometric function. Perception & Psychophysics 1999; 61(1):87–106.

[44] Awiszus F. Fast estimation of transcranial magnetic stimulation motor threshold: is it safe?. Brain Stimulation: Basic, Translational, and Clinical Research in Neuromodulation 2011; 4(1):58–59.

[45] Bergmann TO, Mölle M, Schmidt MA, Lindner C, Marshall L, Born J, Siebner HR. EEG-guided transcranial magnetic stimulation reveals rapid shifts in motor cortical excitability during the human sleep slow oscillation. Journal of Neuroscience 2012; 32(1):243–53.

[46] Zrenner C, Desideri D, Belardinelli P, Ziemann U. Real-time EEG-defined excitability states determine efficacy of TMS-induced plasticity in human motor cortex, Brain Stimulation 2018; 11(2):374–389.

[47] Ding Z, Ouyang G, Chen H, Li X. Closed-loop transcranial magnetic stimulation of real-time EEG based on the AR mode method. Biomed. Phys. Eng. Express 2020; 6:035010.

[48] Pankka H, Roine T, Lehtinen J, Ilmoniemi, RJ. Improving closed-loop TMS timing using the Wavenet model. Brain Stimulation 2021; 14(6):P1636–1637.

[49] Alavi SMM, Goetz SM, Peterchev AV. Optimal estimation of neural recruitment curves using Fisher information: application to transcranial magnetic stimulation. IEEE Transactions on Neural Systems and Rehabilitation Engineering 2019; 27:1320–1330.

[50] Alavi SMM, Goetz SM, Saif M. Input-Output Slope Curve Estimation in Neural Stimulation Based on Optimal Sampling Principles. Journal of Neural Engineering 2021; 18:046071.

[51] Goetz SM, Alavi SMM, Deng Z-D, Peterchev AV. Statistical Model of Motor Evoked Potentials. IEEE Transactions on Neural Systems and Rehabilitation Engineering 2019; 27:1539–1545.

[52] Qi F, Wu AD, Schweighofer N. Fast estimation of transcranial magnetic stimulation motor threshold. Brain Stimul. 2011 Jan;4(1):50–7.

[53] Alavi SMM, Vila-Rodriguez F, Mahdi A, Goetz SM. Closed-loop and automatic tuning of pulse amplitude and width in EMG-guided controllable transcranial magnetic stimulation (cTMS). bioRxiv preprint 2022. doi: 10.1101/2022.05.08.491097

[54] Goetz SM, Luber B, Lisanby SH, Peterchev AV. A Novel Model Incorporating Two Variability Sources for Describing Motor Evoked Potentials. Brain Stimulation 2014; 7:541–552.

[55] Nowak LG, Bullier J. Axons, but not cell bodies, are activated by electrical stimulation in cortical gray matter. I: Evidence from chronaxie measurements. Exp Brain Res. 1998; 118:477–488.

[56] Geddes LA. Accuracy limitations of chronaxie values. IEEE Trans Biomed Eng. 2004; 51:176–181.

